# Cell-Sized Droplet Interfaces Reorganize Protein Secondary Structures through Confinement-Enhanced Membrane Interactions

**DOI:** 10.64898/2026.06.17.732854

**Authors:** Anusuya Pal, Kazutoshi Masuda, Miho Yanagisawa

## Abstract

Cell membranes can regulate protein organization, yet it remains unclear whether membrane-associated structural alterations primarily reflect membrane-induced destabilization or the reorganization of proteins that are already destabilized. How such membrane effects are amplified under cellular-scale confinement also remains poorly understood. Here, we investigate these questions using cell-sized lipid-coated droplets, where the high surface-area-to-volume ratio enhances the contribution of membrane interfaces. Native serum albumin and lysozyme showed little structural reorganization, whereas their thermally denatured forms exhibited pronounced *β*-sheet formation within small droplets when membrane interactions were attractive. For denatured albumin, *β*-sheet-rich organization increased progressively with protein–membrane attraction, while denatured lysozyme selectively formed a localized *β*-sheet-rich shell at a complementary anionic membrane. Circular dichroism spectroscopy independently supported the confinement-enhanced increase in *β*-sheet content of denatured albumin. Fluorescence recovery measurements further revealed strong interfacial arrest in both systems. Together, these results show that membrane interactions can promote structural reorganization of already destabilized proteins through electrostatic recruitment, while cell-sized confinement strongly amplifies the resulting membrane-dependent structural response. Our findings establish cell-sized droplet interfaces as a materials platform for spatially controlling protein structural organization through the interplay of interfacial chemistry and confinement.

## Introduction

Water-in-oil droplets covered by lipid membranes on micrometer-length scales are ubiquitous in pharmaceutical formulations, cosmetics, synthetic cells, and living cells.^1,2^ As the droplet size decreases, the surface-area-to-volume ratio increases, rendering membrane interface effects progressively more dominant. Using droplets covered with lipid membranes as model systems, previous studies have shown that when the droplet radius falls below *∼* 30 *µ*m, the influence of membrane interfaces becomes increasingly pronounced, leading to phenomena such as confinement-enhanced gene expression,^3^ membrane-induced phase separation,^4–6^ and suppressed molecular diffusion.^7–9^ These findings suggest that micrometer-sized droplets, hereafter referred to as cell-sized droplets, are not merely small containers but rather interface-dominated confined systems in which membrane interfaces actively govern molecular organization. Despite increasing recognition of such confinement effects, how interface-dominated confinement influences structural reorganization of biomolecules remains largely unexplored.^10–12^

Proteins provide a prominent example of biomolecules whose structure and assembly can be regulated by interfacial interactions. Membrane interfaces not only recruit proteins but also reshape their conformational landscapes and aggregation pathways.^10,12–14^ For example, membrane interactions have been shown to promote the formation of *β*-sheet-rich amyloid-like structures in a wide range of proteins, including *α*-synuclein, ^11,15,16^ *β*_2_-microglobulin,^17^ and Tau.^18^ A similar role of membranes has been reported in biomolecular condensates formed through liquid–liquid phase separation, where specific lipid–protein interactions promote gelation and amyloid formation of fused in sarcoma (FUS) protein.^19^ These studies demonstrate that membranes strongly influence protein structure and assembly. However, whether these effects arise from membrane-induced destabilization or from the reorganization of proteins that are already destabilized prior to membrane contact remains unresolved.

To address this question experimentally, a model membrane system that can systematically control protein–membrane interactions while avoiding the complexity of the cellular environment is useful. Most previous studies have examined such interactions using planar membranes^10,12,20^ or bulk solutions containing lipid vesicles,^11,18,19^ where confinement effects are negligible. In microscopic compartments, by contrast, the increasing surfacearea-to-volume ratio amplifies the influence of membrane interfaces throughout the confined volume,^4^ raising the possibility that membrane-mediated structural reorganization may become size-dependent. Consistent with this idea, we previously reported that when gelatin, a denatured derivative of collagen, is gelled within lipid-coated droplets, *β*-sheet formation increases as droplet size decreases, resulting in mechanically stiffer microgels.^4,21^ This effect was pronounced in negatively charged membranes, suggesting that membrane electrostatics and confinement act cooperatively to promote structural reorganization. Nevertheless, the respective contributions of the protein conformational state, membrane electrostatics, and confinement-enhanced interfacial dominance remain unclear.^11,19,21^

Here, we use cell-sized lipid-coated droplets as model confined compartments to investigate how membrane interactions reorganize protein structure under interface-dominated confinement. By independently controlling the initial conformational state of proteins, membrane charge, and droplet size, we examine how these factors govern *β*-sheet-rich structural organization in BSA and lysozyme. We show that proteins that are already destabilized prior to membrane contact undergo pronounced membrane-dependent structural reorganization, whereas native proteins exhibit only weak responses. The extent and spatial architecture of this reorganization depend on both membrane electrostatics and protein identity, with confinement amplifying the membrane-dependent response as the droplet size decreases. These findings reveal how interfacial chemistry and cellular-scale confinement cooperate to regulate protein structural organization and provide a controllable materials platform for studying protein structure at membrane interfaces.

## Results

### Structural alteration of proteins inside interface-dominated confinement spaces

To investigate how membrane interactions and micro-confinement cooperatively regulate protein structural alterations, we established a model system in which the protein conformational state, membrane charge, and confinement size could be independently controlled. Cell-sized droplets (with radii *R*_0_ = 5–50 *µ*m) were stabilized by zwitterionic (DOPC), anionic (DOPG), or cationic (DOTAP) lipid monolayers and used as confined environments with distinct interfacial electrostatic properties (Figure 1). As model proteins, we selected bovine serum albumin (BSA; 64 kDa, p*I ≈* 4.7) and lysozyme (Lyz; 14.3 kDa, p*I ≈* 11), which have opposite net charges under neutral conditions. Both native and thermally denatured proteins were encapsulated within droplets, allowing us to distinguish membrane-induced destabilization from membrane-mediated reorganization of already destabilized proteins (see the “Sample Preparation” and “Materials” subsections in the Supporting Information, SI).

**Figure 1:**
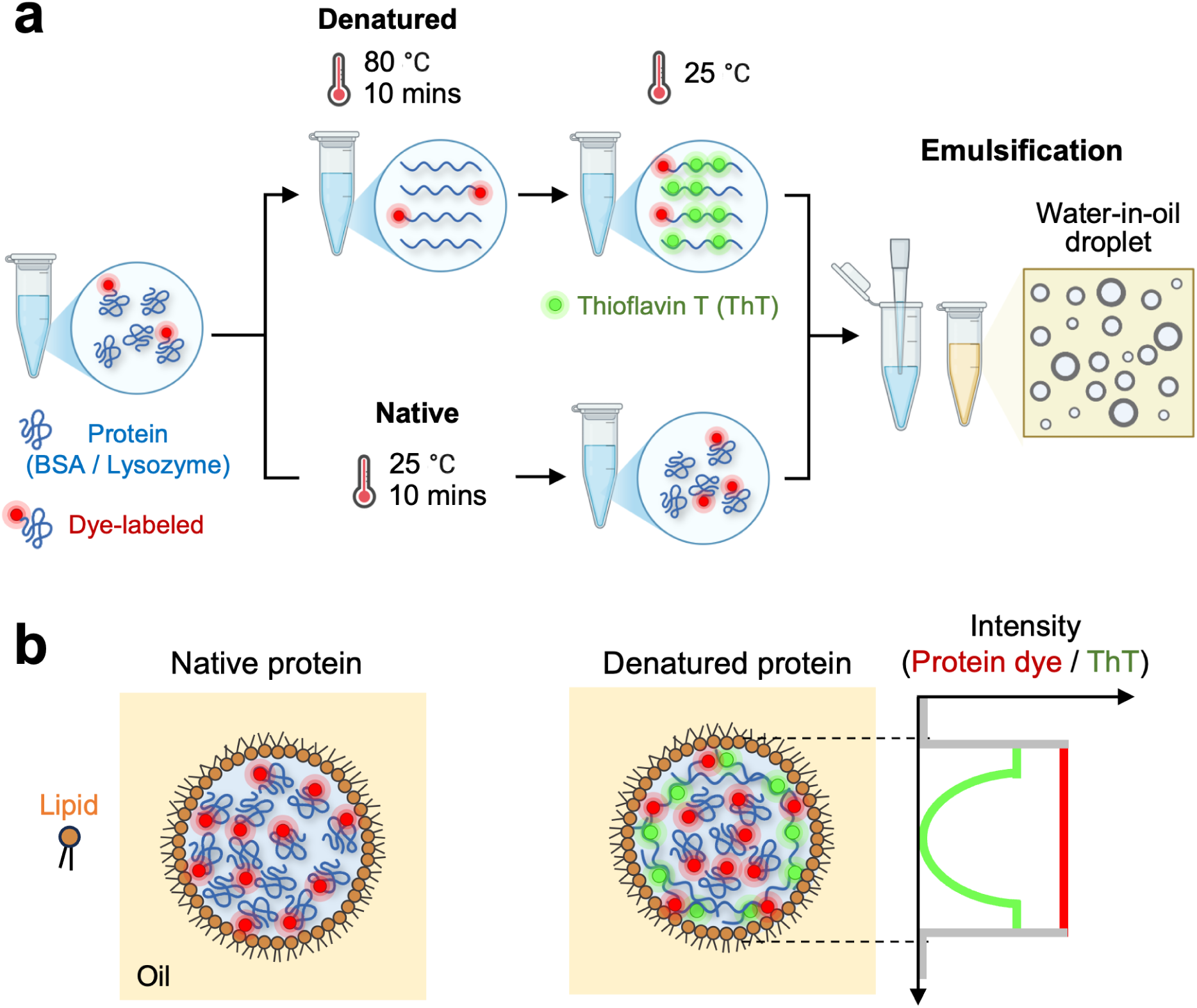
Distinct interfacial behaviors of native and denatured proteins encapsulated in lipid-coated droplets. (a) Native and heat-denatured proteins (blue; bovine serum albumin (BSA) and lysozyme) were encapsulated in water-in-oil droplets covered with a lipid layer. Before emulsification at room temperature (*∼*25 *^◦^*C), protein solutions were heated in bulk at 80 *^◦^*C for 10 min, allowing them to encounter the membrane in a denatured state. (b) Schematics of representative protein redistribution profiles for native proteins (left) and denatured proteins (right). Fluorescence imaging was employed to visualize protein distribution and *β*-sheet-rich structures by using fluorescence-labeled proteins (red) and thioflavin T (ThT, green), respectively.

We combined two complementary fluorescence probes to characterize confinement-induced structural alterations. Thioflavin T (ThT; green) was used to visualize *β*-sheet-rich structures,^22,23^ whereas fluorescently labeled proteins (Texas Red and Rhodamine B; red) were used to characterize protein distribution. Bulk protein solutions without membrane confinement served as reference states, and the fluorescence intensities measured in droplets (*I*) were compared with those in bulk solution (*I*_bulk_).

### Structural change of BSA inside lipid-coated droplets

We first examined how the initial conformational state of BSA influences membrane-mediated structural reorganization in cell-sized droplets. Native BSA exhibited only weak ThT fluorescence irrespective of membrane charge, indicating that confinement alone does not induce substantial *β*-sheet formation (Figure 2a-c, left; anionic DOPG [L(*−*)], zwitterionic

**Figure 2:**
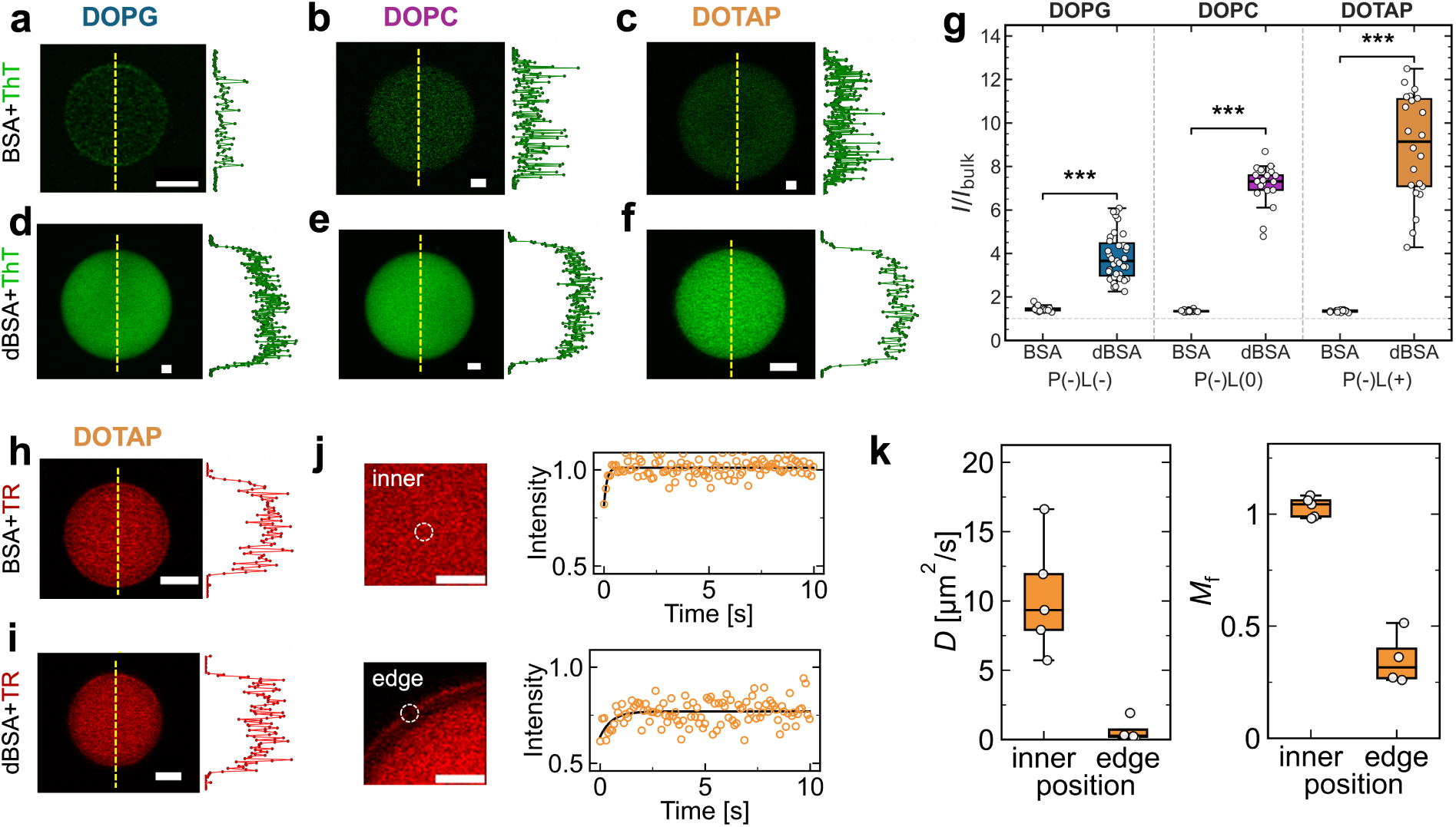
Structural change of BSA in cell-sized droplets. Representative confocal images of native BSA in water-in-oil droplets stabilized by DOPG (a), DOPC (b), and DOTAP (c), imaged using ThT, along with their fluorescence intensity profiles along the dashed center line. (d-f) Corresponding images of denatured BSA (dBSA). (g) Distribution of the confinement enhancement ratio *I/I*_bulk_. Labels P(*−*)L(*−*), P(*−*)L(0), and P(*−*)L(+) denote the charge combinations of protein (P) and lipid (L), respectively. Statistical comparisons between native and denatured conditions (*n ≈* 20) within each lipid were performed using the Mann-Whitney test (p *<* 0.001*^∗∗∗^*). (h-i) Confocal images of BSA and dBSA droplets in DOTAP with dye-labeled proteins, along with their fluorescence intensity profiles along the dashed center line. (j-k) Representative FRAP curves of dBSA in DOTAP droplets (inner and edge parts, indicated by dashed white circles in the images (j)) and the values of the diffusion coefficient (*D*) and mobile fraction (*M*_f_) obtained from the analysis. All scale bars are 10 *µ*m.

DOPC [L(0)], and cationic DOTAP [L(+)]). The fluorescence remained uniformly distributed throughout the droplet interior, and the corresponding intensity profiles (Figure 2a-c; right) showed no evidence of localized interfacial enrichment. Thus, although BSA experiences confinement within cell-sized droplets, its native structure appears largely resistant to membrane-mediated structural conversion.

In contrast, thermally denatured BSA (dBSA) displayed a striking increase in ThT fluorescence under all membrane conditions (Figure 2d-f), demonstrating that the susceptibility of the protein to structural reorganization is strongly dependent on its initial conformational state. Notably, the enhanced ThT fluorescence remained spatially distributed throughout the droplet rather than forming a localized interfacial layer. This suggests that membrane interactions within a cell-sized space facilitate structural transformation without substantial condensation on the membrane.

Structural reorganization was quantified using the normalized ThT fluorescence intensity ratio (*I/I*_bulk_), where *I* and *I*_bulk_ denote the mean fluorescence intensities in droplets and bulk solution, respectively. Native BSA remains close to the bulk reference level under all lipid conditions (*I/I*_bulk_ = 1.43 for DOPG, 1.35 for DOPC, and 1.36 for DOTAP; Figure 2g). Here, P(*−*)L(*−*), P(*−*)L(0), and P(*−*)L(+) denote negatively charged protein (P) paired with negatively charged, neutral, and positively charged lipids (L), respectively. In contrast, dBSA exhibits substantially elevated values that increase monotonically with membrane charge, from 3.66 in DOPG [L(*−*)], to 7.31 in DOPC [L(0)], and 9.14 in DOTAP [L(+)]. This enhancement observed upon denaturation was significant under all lipid conditions (Mann–Whitney test with Holm–Bonferroni correction; *p <* 0.05*^∗^*; Table ST1 in Supporting Information (SI)). A two-way ANOVA further revealed a strong protein state *×* lipid interaction (*F* (2, 122) = 54, *p <* 0.001, *η*^2^ = 0.47; Table ST2 in SI), indicating that the influence of membrane composition depends strongly on the conformational state of the protein. Moreover, all pairwise comparisons among the dBSA conditions were significant, confirming the ordering DOPG *<* DOPC *<* DOTAP (Figure S1 and Table ST3 in SI). Because BSA carries a net negative charge at neutral pH conditions, this ordering indicates that increasingly attractive protein–membrane electrostatic interactions enhance confinement-induced *β*-sheet formation.

To assess whether the enhanced ThT fluorescence arises from protein accumulation or structural reorganization, we next visualized the total protein distribution using dye-labeled BSA. Both native BSA and dBSA exhibited spatially uniform fluorescence throughout DOTAP droplets, with no detectable enrichment at the interface (Figure 2h-i). Although the fluorescence intensity of dye-labeled BSA was approximately twofold higher in droplets than in bulk solution, likely reflecting partial adsorption of proteins to the glass surface in the bulk reference measurements, no significant dependence on lipid composition was observed (Figures S2, S3 in SI). Together with the uniformly distributed ThT signal observed for dBSA (Figure 2d-f), these results indicate that the enhanced ThT fluorescence arises from spatially distributed *β*-sheet formation rather than preferential protein accumulation on the membrane or the formation of a templated interfacial shell.

We characterized protein dynamics at the membrane interface using fluorescence recovery after photobleaching (FRAP) measurements (see the FRAP subsection in SI). FRAP measurements performed on dBSA in DOTAP droplets revealed markedly slower recovery at the interface than in the droplet interior (Figure 2j). Whereas proteins in the interior remained highly mobile (with diffusion coefficients of *D ≈* 5–17 *µ*m^2^s*^−^*^1^ and mobile fractions *M*_f_ *≈* 1.0), the interfacial population exhibited strongly suppressed mobility and a reduced mobile fraction (*D ≈* 0 *µ*m^2^s*^−^*^1^, *M*_f_ *≈* 0.3; Figure 2k). Thus, despite the absence of detectable protein enrichment (Figure 2i), dBSA undergoes dynamic arrest at the membrane interface. These observations suggest that attractive protein–membrane interactions promote structural reorganization through interfacial trapping, while the resulting *β*-sheet-rich structures remain distributed throughout the confined space (Figure 2f).

### Structural change of lysozyme inside lipid-coated droplets

We further examined the generality of confinement-enhanced structural reorganization using lysozyme, a protein with a net charge opposite to that of BSA under neutral conditions. As in BSA [P(*−*)], native lysozyme (Lyz, [P(+)]) exhibits only weak ThT fluorescence under all membrane conditions (Figure 3a-c; From left to right: anionic DOPG [L(*−*)], zwitterionic DOPC [L(0)], and cationic DOTAP [L(+)]). It indicates that confinement alone is insufficient to induce substantial *β*-sheet formation. In contrast, thermally denatured lysozyme (dLyz) underwent pronounced structural reorganization (Figure 3d-f), demonstrating that susceptibility to membrane-mediated structural conversion again depends strongly on the protein’s initial conformational state.

**Figure 3:**
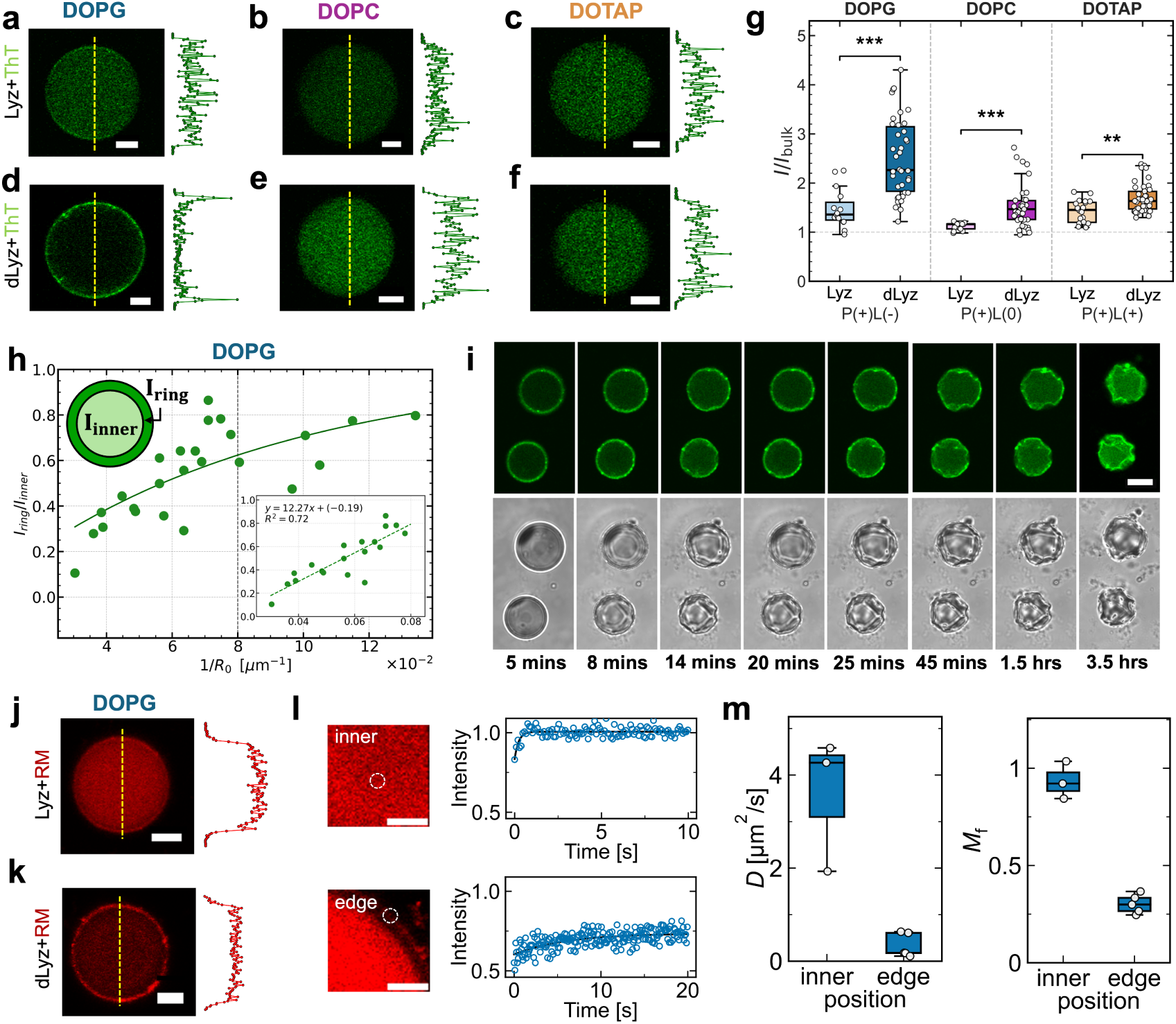
Structural change of lysozyme in cell-sized droplets. Representative confocal images of native Lyz in droplets stabilized by DOPG (a), DOPC (b), and DOTAP (c), imaged using ThT, along with their fluorescence intensity profiles along the dashed center line. (d-f) Corresponding images of denatured Lyz (dLyz). (g) Distribution of the confinement enhancement ratio *I/I*_bulk_. Labels P(+)L(*−*), P(+)L(0), and P(+)L(+) denote the charge combinations of protein (P) and lipid (L), respectively. Statistical comparisons between native and denatured conditions (*n ≈* 20) within each lipid were performed using the Mann-Whitney test (p *<* 0.01*^∗∗^* and p *<* 0.001*^∗∗∗^*). (h) Dependence of the intensity ratio (*I*_ring_*/I*_inner_) on inverse droplet radius (1*/R*_0_) for dLyz/DOPG droplets. (i) Time evolution of the dLyz/DOPG interfacial ring monitored by ThT (top) and bright-field imaging (bottom), showing progressive shell-like interfacial organization and subsequent buckling over time. (j-k) Confocal images of Lyz and dLyz in DOPG droplets with dye-labeled proteins, along with their fluorescence intensity profiles along the dashed center line. (l-m) Representative FRAP curves of dLyz in DOPG droplets (inner and edge parts) and the values of the diffusion coefficient (*D*) and mobile fraction (*M*_f_) obtained from the analysis. Photobleached regions are indicated by dashed white circles in the (l) images. All scale bars are 10 *µ*m.

Unlike dBSA, however, dLyz [P(+)] displayed a striking dependence on membrane identity. Under the complementary electrostatic condition provided by anionic DOPG [L(*−*)], ThT fluorescence became strongly localized at the droplet periphery, producing a pronounced ring-like structure accompanied by reduced fluorescence in the droplet interior (Figure 3d). In contrast, dLyz remained uniformly distributed in DOPC and DOTAP droplets without detectable interfacial enrichment (Figure 3e-f). Thus, whereas dBSA generated spatially homogeneous *β*-sheet-rich structures, dLyz formed a distinct interfacial assembly selectively under attractive protein–membrane interactions.

These differences were reflected in the normalized ThT fluorescence intensity (*I/I*_bulk_) as shown in Figure 3g. Native Lyz remained close to the bulk reference under all lipid conditions (1.36 for DOPG, 1.16 for DOPC, and 1.45 for DOTAP). By contrast, dLyz exhibited its strongest response in DOPG (*I/I*_bulk_ = 2.26), whereas DOPC and DOTAP produced only modest enhancements (1.47 and 1.63, respectively). The enhancement observed upon denaturation was significant under all lipid conditions (Mann–Whitney test, *p <* 0.05*^∗^*; see also Table ST4 in SI). A two-way ANOVA further revealed the significant protein state *×* lipid interaction (*F* (2, 158) = 6.49, *p* = 0.002, *η*^2^ = 0.08; Table ST5 in SI), confirming that membrane composition influences structural reorganization differently in native and denatured lysozyme. Therefore, unlike dBSA, which displayed a monotonic dependence on membrane charge, dLyz responds selectively to the complementary DOPG membrane (see also Table ST6 and Figure S4 in SI).

To characterize the droplet size dependence of the shell-like structure, we quantified the relative contribution of the ring structure using the ratio (*I*_ring_*/I*_inner_), defined as (*I*_whole_ *− I*_inner_)*/I*_inner_, where *I*_whole_ is the mean fluorescence intensity within the whole droplet and *I*_inner_ is the mean intensity within the center region (see the Image anaLyzis subsection in SI). In dLyz/DOPG droplets, the ring contribution increased systematically with decreasing droplet size (1*/R*_0_ *>* 0.03 *µ*m*^−^*^1^ (radii *R*_0_ *<* 33 *µ*m), Figure 3h), indicating that confinement progressively enhances interfacial structural organization. As shown in the inset, *I*_ring_*/I*_inner_ scales approximately linearly with 1*/R*_0_ (*I*_ring_*/I*_inner_ = 12.27 (1*/R*_0_) *−* 0.19, *R*^2^ = 0.72). At smaller droplets (*R*_0_ *<* 10 *µ*m), shell-like structures were accompanied by buckling of the droplets (Figure 3i). The deviation from the spherical shape can be seen within roughly 20 minutes of droplet formation, and this buckling process persists for 3.5 hours. Because Laplace pressure tends to maintain a spherical shape, buckling cannot occur in purely viscous droplets and instead requires a finite elastic nature capable of supporting compressive stresses. Thus, the emergence of buckling demonstrates that the interfacial *β*-sheet-rich assembly behaves not only as a dynamically arrested layer but also as an elastic shell. Neither size-dependent shell enhancement nor buckling was observed in dBSA droplets under any membrane conditions within the investigated size range.

The total protein distribution was visualized using dye-labeled Lyz to clarify whether the interfacial shell reflects protein accumulation or localized structural reorganization (Figure 3j-k, left). In contrast to the pronounced ThT ring observed in dLyz/DOPG droplets, the protein fluorescence (red) remains uniformly distributed throughout the droplet interior without significant interfacial enrichment, as confirmed by the corresponding intensity profiles (Figure 3j-k, right). Consistent with this observation, the value of *I*_ring_*/I*_inner_ for dye-labeled Lyz is negative, whereas it was strongly positive for Tht (Figure S5 in the SI).

Additionally, the normalized fluorescence intensity of dye-labeled Lyz relative to the bulk remained close to unity and showed no significant dependence on lipid composition (Figures S6, S7 in the SI). Moreover, dBSA/DOTAP droplets exhibited a higher value of *I/I*_bulk_ than dLyz/DOPG droplets despite lacking an interfacial shell (Figure S8 in SI), suggesting that shell formation does not correlate with the total amount of protein present within droplets. These results demonstrate that the ThT-positive shell in dLyz/DOPG droplets does not arise from local protein accumulation but from localized formation of *β*-sheet-rich structures at the membrane interface.

The dynamics of the dLyz interfacial shell were examined by FRAP using dye-labeled lysozyme (Figure 3l-m). Similar to dBSA/DOTAP droplets, proteins in the droplet interior remained highly mobile (*D ≈* 2–5 *µ*m^2^s*^−^*^1^ and *M*_f_ *∼* 1), whereas the interfacial population exhibited strongly suppressed mobility and a reduced mobile fraction (*D ≈* 0 *µ*m^2^s*^−^*^1^ and *M*_f_ *∼* 0.3). Therefore, despite their distinct structural organizations (spatially homogeneous *β*-sheet-rich assemblies for dBSA/DOTAP; Figure 2f and interfacial shell structures for dLyz/DOPG; Figure 3d), both systems exhibit pronounced dynamic arrest at the membrane interface. Notably, the *D* within the interior of dLyz/DOPG is approximately 3*×* lower than that of dBSA/DOTAP, indicating reduced mobility of the lysozyme population throughout the confined volume, relative to dBSA.

### Homologous proteins exhibit similar membrane-dependent structural architectures

To examine whether the observed structural changes extend beyond the specific protein sequences studied here, we tested two homologous proteins under the lipid conditions that produced the strongest responses in each system. Despite differences in amino acid sequence, both homologs exhibited structural architectures similar to those observed for the corresponding proteins. Denatured human serum albumin (dHSA) exhibited spatially homogeneous *β*-sheet-rich fluorescence in DOTAP droplets, analogous to dBSA (Figure S9 in SI).

Denatured human lysozyme (dhLyz) formed a pronounced interfacial shell in DOPG droplets, and small droplets with *R*_0_ *<* 10 *µ*m underwent buckling, similar to dLyz (Figure S10 in SI). These observations suggest that the confinement-enhanced structural reorganization driven by attractive protein–membrane interactions is not limited to the specific protein sequences examined here. The preservation of similar structural architectures in these homologs further suggests that protein-level physicochemical properties, rather than sequence-specific features alone, may contribute to the membrane-mediated structural response.

### CD spectra of BSA in bulk and in droplets

To further support the conformational changes suggested by the ThT experiments, we performed circular dichroism (CD) measurements on native and denatured BSA in bulk solution and in DOTAP droplets. For BSA in droplets, we centrifuged the emulsion at 7000 *×* g for 5 min and collected the aqueous phase for CD measurements. Because Lyz in droplets was strongly localized on the membrane and could not be quantitatively collected by centrifugation, we used BSA as a representative protein for the CD measurements. The CD spectrum of native BSA in bulk solution agreed well with its expected secondary structure and with previous reports in the literature.^24^ To quantify the secondary-structure contributions to the CD spectra, the spectra were analyzed using the BeStSel web server.^25^ The estimated secondary-structure compositions of native BSA in bulk solution and in droplets were nearly identical (Figure 4, left), with an *α*-helix content of approximately 62% and essentially no detectable *β*-sheet contribution, consistent with previous reports and the expected secondary structure of native BSA.^24,26^ In contrast, thermal denaturation induced substantial changes in the estimated secondary-structure composition of BSA. Moreover, we observed a pronounced difference between denatured BSA in bulk solution and in droplets (Figure 4, right). The *α*-helix contribution decreased from 34.9% in bulk solution to 21.2 % in droplets, whereas the *β*-sheet contribution increased from 11.6% to 18.9%. These results are consistent with the ThT measurements and support a scenario in which, after partial thermal denaturation, attractive membrane interactions under confinement further promote the formation of *β*-sheet-rich structures in BSA.

**Figure 4:**
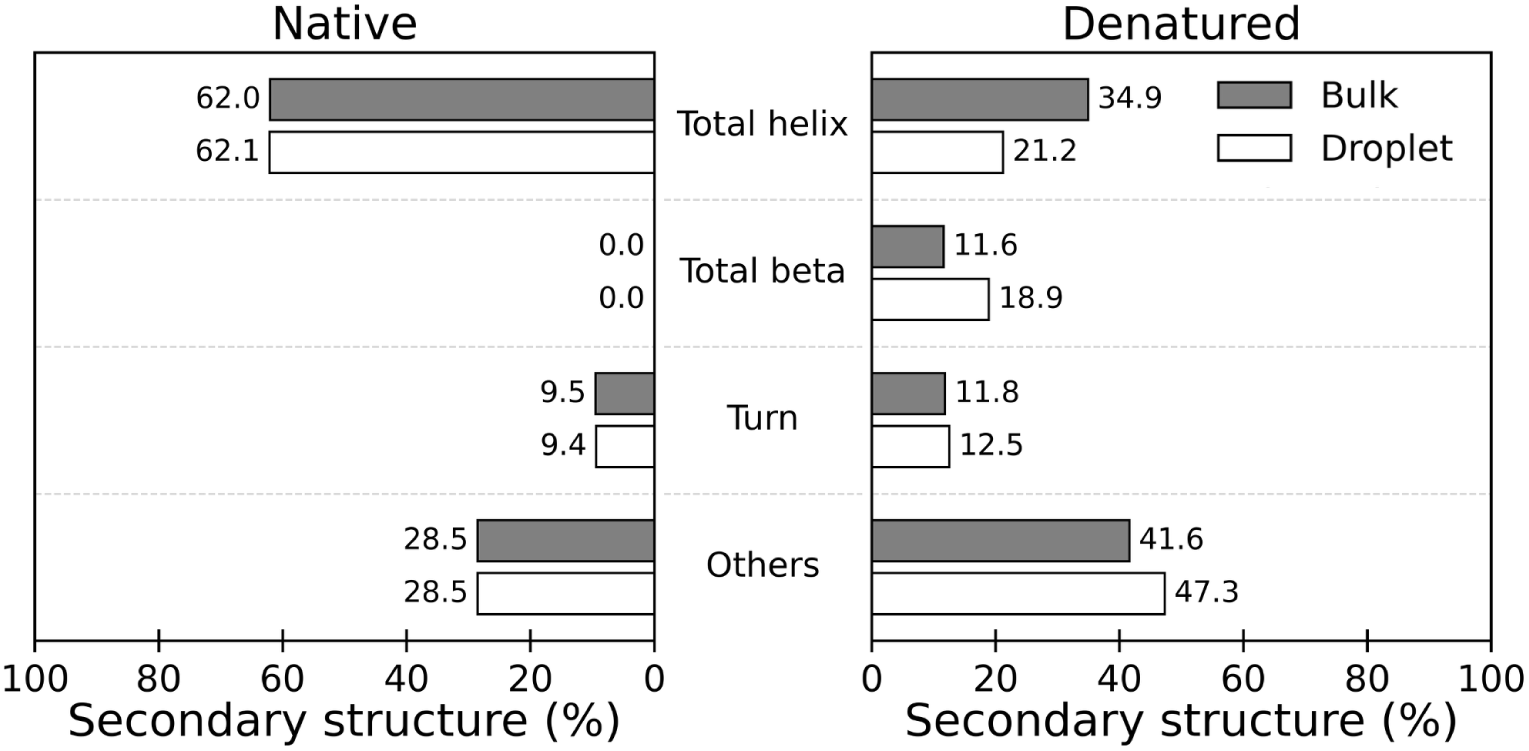
Estimated secondary structure composition from circular dichroism (CD) spectra. The secondary structure compositions of native and denatured BSA in bulk solution and in DOTAP droplets were estimated using the BeStSel web server. Native BSA is shown on the left, and denatured BSA on the right.

## Discussion

### Confinement and denaturation promote membrane-mediated *β*-sheet reorganization

The central finding of this study is that pronounced membrane-mediated *β*-sheet reorganization occurs when protein destabilization is combined with cell-sized confinement, as illustrated in Figure 5(i). Native BSA and lysozyme exhibited only weak structural responses regardless of membrane composition, whereas denatured proteins underwent pronounced membrane-dependent reorganization. Because denaturation was induced in bulk before membrane contact, the observed structural reorganization is likely originates not from membrane-induced denaturation itself, but rather from subsequent interactions between the destabilized protein and the membrane. Thermal denaturation may partially unfold the proteins, exposing charged or otherwise membrane-interacting regions and facilitating membrane engagement. These observations are consistent with previous studies showing that membrane interfaces can regulate protein aggregation pathways by stabilizing assembly-prone conformations.^15,27,28^

**Figure 5:**
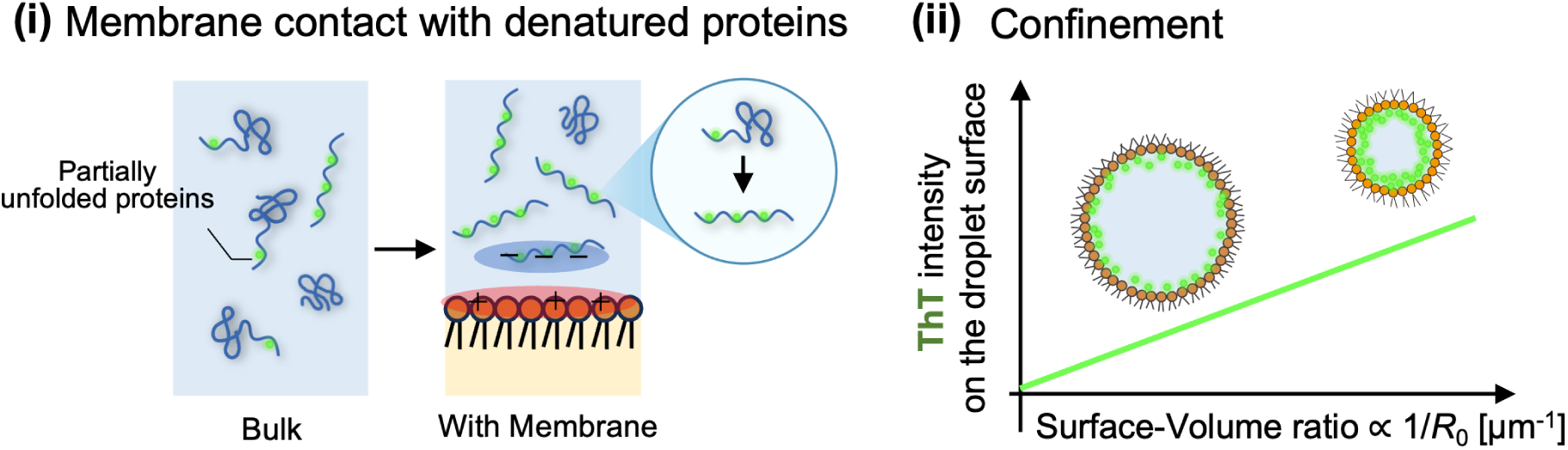
Two primary factors in membrane-induced structure transition. (i) Proteins are partially unfolded in the bulk solution by heating. Partial unfolding exposes charged segments, thereby enhancing protein–membrane contact and increasing the affinity of proteins for the membrane (orange). The formation of *β*-sheet-rich structures is detected by ThT (green). (ii) Confinement further amplifies membrane effects, as evidenced by the droplet-size dependence of ThT intensity at the droplet surface, with smaller droplets displaying stronger interfacial ThT signals.

Importantly, the CD measurements independently support this interpretation. Whereas native BSA showed nearly identical secondary-structure compositions in bulk solution and droplets, denatured BSA exhibited a further decrease in *α*-helix content and an increase in *β*-sheet content when confined within DOTAP droplets (Figure 4). The increase in *β*-sheet content from 11.6 % in bulk to 18.9 % in droplets is consistent with the enhanced ThT fluorescence observed under confinement, supporting the interpretation that membrane interactions under confinement promote further structural reorganization of already denatured BSA. Although we performed CD measurements only for BSA, agreement between the CD and ThT measurements strengthens the assignment of the fluorescence response to changes in protein secondary structure rather than to changes in protein concentration alone.

Our results further indicate that cell-sized confinement amplifies membrane-mediated structural reorganization (Figure 5(ii)). Even under membrane conditions that did not produce pronounced shell formation, ThT fluorescence remained elevated relative to bulk solution (see dBSA case in Figure 2). Similar confinement-dependent phenomena have been reported in lipid-coated droplets, including membrane-regulated gene expression and confinement-induced *β*-sheet transitions of gelatin.^3,21^ As the droplet size decreases, the increasing surface-area-to-volume ratio enhances the influence of the membrane interface on the confined volume. Within this framework, confinement does not determine the specific architecture that emerges; rather, it increases the extent to which membrane-dependent structural reorganization can occur.

An important implication is that the observed structures cannot be explained solely by increased protein concentration. Although dye-labeled proteins exhibited enhanced droplet-to-bulk fluorescence under multiple conditions (Figures S3, S7 in SI), their spatial distributions within droplets remained homogeneous and did not show lipid dependence. In particular, the ThT-positive shell observed for dLyz/DOPG was not accompanied by detectable protein enrichment at the membrane interface (Figure S5 in SI). Thus, membrane-mediated structural reorganization reflects a localized change in protein organization rather than a simple accumulation of protein mass.

### A two-step mechanism links electrostatic recruitment to protein-dependent structural organization

The present results support a hierarchical mechanism in which membrane interactions govern interfacial engagement, whereas protein-specific properties influence the structural architecture that develops following membrane recruitment. Despite their markedly different morphologies, both dBSA/DOTAP and dLyz/DOPG exhibit strong dynamic arrest at the membrane interface, with substantially reduced mobility relative to the droplet interior (Figures 2j-k and 3l-m). Thus, interfacial trapping may represent a common intermediate in the membrane-mediated structural reorganization observed here. The electrostatic basis of this membrane engagement is supported by the charge states of the proteins and lipids: pH measurements of the protein solutions confirmed that they were above the albumin pI for BSA/HSA and well below the lysozyme pI for Lyz/hLyz in both native and denatured states (Table ST7 in SI). These charge relationships are consistent with preferential interactions between oppositely charged proteins and lipid membranes.

The architectural divergence emerges after membrane recruitment. Under strongly engaging conditions, dBSA exhibited spatially homogeneous *β*-sheet-associated ThT fluorescence throughout the DOTAP droplet interior, whereas dLyz forms a localized shell-like structure at the DOPG membrane. These contrasting outcomes suggest that the membrane interface alone does not dictate the final structure. Rather, membrane interactions determine the extent and location of protein–membrane engagement, while intrinsic protein properties influence how structural reorganization proceeds after recruitment. Similar membrane-dependent aggregation pathways have been reported for amyloid-forming proteins, including *α*-synuclein and tau, in which membrane interfaces strongly influence the pathways leading to distinct protein assemblies.^12,13,15^

One possible origin of this protein dependence is disulfide topology. dBSA contains seventeen intrachain disulfides distributed across predominantly *α*-helical domains, ^29,30^ whereas lysozyme contains only four disulfides and retains a comparatively extended region associated with its *β*-domain.^31,32^ Following thermal denaturation, these differences in disulfide connectivity and domain organization may influence the conformational flexibility and accessibility of *β*-prone regions, thereby contributing to the distinct structural architectures observed here. Consistent with this possibility, the lower diffusion coefficient observed within dLyz/DOPG droplets relative to dBSA/DOTAP suggests stronger protein association in the lysozyme system, in agreement with the known propensity of unfolded lysozyme to form self-associated states.^33,34^ Moreover, similar structural architectures were also observed for the corresponding human homologs (Figures S9 and S10 in SI), suggesting that the responses are not strictly sequence-specific. These observations indicate that protein-level physicochemical properties can contribute to the structural outcome following membrane recruitment, although the molecular determinants of this protein dependence remain to be established.

## Conclusion

Together, our results show that cell-sized confinement can amplify membrane-mediated structural reorganization of proteins that are already destabilized. Electrostatic interactions promote protein–membrane engagement, while intrinsic protein properties influence the resulting structural architecture, leading to distinct spatial organizations for BSA and lysozyme. The observation of similar architectures in the corresponding human homologs further suggests that these responses are not strictly sequence-specific. This framework provides a physical basis for understanding how similar membrane environments can generate distinct protein structural organizations and may provide insight into how membrane interfaces influence the formation of amyloid-associated structures in biological and materials contexts.^17,28,35^

## Data availability

The data supporting the findings of this study are available within the article and its Supporting Information. Additional datasets, raw image files, and analysis scripts are available from the corresponding author upon reasonable request.

## Supporting Information

The Supporting Information is available free of charge at [URL]. Section S1: Experimental details; Section S2: Statistical analyses of fluorescence images of BSA samples; Table ST1: Native versus denatured BSA; Table ST2: Two-way ANOVA for BSA; Table ST3: Pairwise comparisons for denatured BSA; Figure S1: Lipid-dependent ThT fluorescence for denatured BSA in droplets; Figure S2: Fluorescence intensity profiles of dye-labeled native BSA along the droplet center; Figure S3: Relative fluorescence intensity of dye-labeled native BSA in droplets compared with the bulk solution; Section S3: Statistical analyses of fluorescence images of lysozyme samples; Table ST4: Native versus denatured lysozyme; Table ST5: Two-way ANOVA for lysozyme; Table ST6: Pairwise comparisons for denatured lysozyme; Figure S4: Fluorescence intensity profiles of dye-labeled native lysozyme along the droplet center; Figure S5: Protein distribution in denatured lysozyme in DOPG droplets; Figure S6: Fluorescence intensity profiles of dye-labeled native lysozyme; Figure S7: Relative fluorescence intensity of dye-labeled native lysozyme in droplets compared with the bulk solution; Figure S8: Relative fluorescence intensity of dye-labeled denatured proteins in droplets compared with the bulk solution; Figure S9: Fluorescence intensity profiles of ThT for denatured HSA along the DOTAP droplet center; Figure S10: Buckled DOPG droplets with denatured human lysozyme.

## Acknowledgement

We thank Hiroshi Masai and Yutaro Kawano from The University of Tokyo for their experimental support with CD analysis. This research was partially funded by the Japan Society for the Promotion of Science (JSPS) KAKENHI (grant nos. 22H01188, 24H02287 (M. Y.); 26K17105 (A. P.)), the Japan Science and Technology Agency (JST) (grant nos. FOREST (JPMJFR213Y), CREST (JPMJCR22E1) (M. Y.)), and the World-Leading Innovative Graduate Study Program for Advanced Basic Science Course (WINGS-ABC) at the University of Tokyo (K. M.).

