## Supplemental information file for "Cell-Sized Droplet Interfaces Reorganize Protein Secondary Structures through Confinement-Enhanced Membrane Interactions"

### S1. Experimental

#### Materials

Bovine serum albumin (BSA; ≥98%, fatty-acid free; catalog no: A6003-5G), human serum albumin (HSA; ≥96%; catalog no: A1653-5G), recombinant human lysozyme (≥95%; catalog no: L1667-1G), and thioflavin T (ThT; catalog no: T3516-5G) were obtained from Sigma–Aldrich (USA). The hen egg-white lysozyme (catalog no: 100831) was purchased from MP Biomedicals, USA. ThT, Fluorescence-labeled BSA (Texas-Red (TR)-BSA; catalog no: A2307; Invitrogen, CA, USA) and fluorescence-labeled lysozyme (Rhodamine-B (RB)-Lys; catalog no: LS1-RB-1; Nanocs, USA) were used for fluorescence imaging experiments. 1,2-dioleoyl-*sn*-glycero-3-phosphocholine (DOPC; catalog no: 850375C), 1,2-dioleoyl-*sn*-glycero-3-phospho-(1′-*rac*-glycerol) sodium salt (DOPG; catalog no: 840475C), and 1,2-dioleoyl-3-trimethylammonium-propane chloride (DOTAP; catalog no: 890890P) were obtained from Avanti Polar Lipids (USA). Hexadecane (≥99%, anhydrous; catalog no: 07819-32; Nacalai Tesque, Kyoto, Japan) was used as the oil phase. Milli-Q water (18.2 MΩ cm) was used throughout. All materials were used as received.

#### Sample preparation

Aqueous protein solutions were prepared at 5 wt% using Milli-Q water. ThT and dye-labeled proteins were also prepared in Milli-Q water. Protein solutions containing the dye-labeled proteins were placed in sealed Eppendorf tubes and heated at 80 ℃ for 10 min to obtain denatured proteins. After thermal treatment, the samples were cooled to 25 ℃ using a temperature-controlled block before adding ThT. Because denatured protein solutions can transition into a gel-like state upon prolonged standing, all denatured samples were freshly prepared. For fluorescence imaging, we added ThT and dye-labeled proteins to the protein samples to achieve a final concentration of 5 mM for each component.

Lipid stock solutions were first prepared in chloroform at 10 mM. Aliquots were transferred into glass tubes, and the solvent was evaporated under a gentle nitrogen stream to form dry lipid films. Hexadecane was subsequently added to obtain a final lipid concentration of 2 mM. The lipid/oil mixtures were sonicated at 60 ℃ for 90 min and equilibrated at room temperature (approximately 25 ℃) to ensure homogeneous lipid dissolution prior to droplet preparation.

Water-in-oil droplets encapsulated by lipid monolayers were prepared by adding 2 µL of the aqueous protein-dye solution to 40 µL of the lipid-containing hexadecane phase as reported previously.[S1] The mixtures were equilibrated for 5 min to enable lipid adsorption at the nascent water-oil interface before droplet formation. Droplets were then generated by gentle manual agitation (tapping), just enough to form droplets without causing extensive bulk emulsification. The resulting droplet radius ($R_0$) ranges from approximately 1 to 50 µm. However, to eliminate the influence of droplet curvature in fluorescence intensity analysis, only droplets with $R_0 > 5$ µm were used in the analysis. Aliquots were placed into glass-bottom imaging dishes coated with the lipid-oil phase to reduce droplet adhesion, and imaging was performed within 30 min of preparation.

#### pH measurements

The pH of 5 wt% aqueous protein solutions was measured for both native and denatured states using a HORIBA LAQUA 9618N pH electrode connected to a HORIBA LAQUAact D-71 pH meter. The electrode was calibrated prior to measurement using HORIBA standard pH buffer solutions at pH 4 (model 100-4), pH 7 (model 100-7), and pH 9 (model 100-9). Each sample was measured five times, and values are reported as mean ± standard deviation.

#### Confocal microscopy

Confocal images were acquired on a laser-scanning microscope (IX83, FV1200, Olympus, Japan), equipped with a water-immersion objective lens (UPLSAPO 60XW, NA 1.4, Olympus, Japan) with 488 nm excitation for ThT (band-pass 500-540 nm), 561 nm for RB (band-pass 575-625 nm), and 594 nm for TR (band-pass 610-660 nm). For each protein system (BSA, HSA, Lys, and hLys), imaging parameters including high voltage (HV), laser power, detector gain, and scanning rate were maintained constant across all corresponding native, denatured, and lipid conditions to enable quantitative comparison within the same protein background. Because fluorescence response varied between protein systems and fluorophore combinations, acquisition settings were optimized independently for each protein system. Bulk reference measurements ($I_{\mathrm{bulk}}$) were acquired under the identical optical configuration used for the corresponding confined droplet measurements.

#### FRAP

FRAP experiments for dBSA and dLys were performed using the same confocal microscopy setup as described above. To probe the mobility of proteins, 1/1000 of the total dBSA and dLys molecules were replaced with fluorescently labeled dBSA-TR and dLys-RM, respectively. Circular regions of interest with a radius of $w \approx 2\,\mu$m were photobleached by applying a high-intensity laser pulse. Fluorescence recovery was then monitored over time under the same imaging conditions as before photobleaching. The fluorescence intensity in the bleached region was corrected for background fluorescence and normalized by the pre-bleach intensity. The pre-bleach intensity, $I_{\mathrm{pre}}$, was defined as the average fluorescence intensity immediately before photobleaching. The intensity immediately after photobleaching was denoted as $I_0$, and the plateau intensity after recovery was denoted as $I_\infty$. All recovery profiles were well described by a single-exponential model with a relaxation time $\tau$: $I(t) = I_0 + (I_\infty - I_0)\left(1 - e^{-t/\tau}\right)$. The apparent diffusion coefficient $D$ was estimated from the relaxation time as $D = \frac{w^2}{4\tau}$, where $w$ is the radius of the bleached circular region. This relation provides an approximate estimate of the effective diffusion coefficient from the characteristic recovery time. The mobile fraction $M_{\mathrm{f}}$ was calculated as the fraction of fluorescence recovered relative to the fluorescence lost by photobleaching, as $M_{\mathrm{f}} = \frac{I_\infty - I_0}{I_{\mathrm{pre}} - I_0}$.

#### Image analysis

Image analysis was performed in Fiji.[S2] For each droplet, the droplet boundary was identified from either the transmitted-light or fluorescence channel, and a circular region of interest (ROI) was fitted to the droplet contour. $R_0$ was calculated from the droplet perimeter/$2\pi$. Mean droplet fluorescence intensity ($I$) was obtained using a circular ROI encompassing the full droplet area. The confinement enhancement ratio ($I/I_{\text{bulk}}$) was calculated by normalizing the mean intradroplet intensity to the corresponding unconfined bulk reference intensity ($I_{\text{bulk}}$) acquired from a matched-concentration protein solution under identical imaging conditions.

To quantify interfacial enhancement, the ring intensity ($I_{\text{ring}}$) for each droplet was defined as $I_{\text{ring}} = I_{\text{whole}} - I_{\text{inner}}$, where $I_{\text{whole}}$ denotes the mean fluorescence intensity measured within the full droplet ROI and $I_{\text{inner}}$ denotes the mean intensity within a concentric central ROI. Here, $I_{\text{ring}}$ represents the fluorescence contribution associated with the interfacial region relative to the droplet interior. To account for droplet-to-droplet intensity variation, the ring contribution was normalized by the inner intensity, yielding the dimensionless ratio as $\frac{I_{\text{ring}}}{I_{\text{inner}}} = \frac{I_{\text{whole}} - I_{\text{inner}}}{I_{\text{inner}}}$.

#### Statistics

All analyses were performed in Python using *scipy.stats* and *statsmodels*.[S3,S4] The $I/I_{\text{bulk}}$ ratio was log-transformed prior to two-way ANOVA to satisfy normality assumptions, so pairwise comparisons used non-parametric Mann-Whitney U tests with Holm-Bonferroni correction for multiple comparisons within each protein. Sample sizes (number of droplets per condition) and statistical test outputs are tabulated in the Supporting Information. A significance threshold of $p < 0.05$ was used throughout the study.

#### Circular dichroism (CD) measurements

CD spectroscopy was used to investigate the secondary structure of the proteins. Briefly, we recorded CD spectra on a Jasco J−1100 spectropolarimeter (Jasco Inc., Japan) using a quartz cuvette with a 1 mm light path. We acquired spectra from 260 to 190 nm at a resolution of 0.1−100 nm and 25 °C. Final spectra represent the average of five accumulated scans. We prepared a native and denatured 5 wt% (750 $\mu$M BSA stock solution and diluted it to 30 $\mu$M. We collected the confined BSA solutions inside droplets by centrifugation at 25 °C at 7000 × g for 5 min. Acquired spectra were converted to molar ellipticity by the equation $[\theta] = (\theta)/(10 \times c \times l \times nr)$, where $[\theta]$ is the molar ellipticity ($\mathrm{deg, cm^2, dmol^{-1}}$), $(\theta)$ is the experimentally observed ellipticity ($\mathrm{deg, cm^2, dmol^{-1}}$), $l$ is the path length (mm; $l = 10$ for this experiment), and $nr$ is the number of residues in the peptide ($nr = 583$ for BSA).

(S3) Virtanen, P.; Gommers, R.; Oliphant, T. E.; Haberland, M.; Reddy, T.; Cournapeau, D.; Burovski, E.; Peterson, P.; Weckesser, W.; Bright, J.; van der Walt, S. J.; Brett, M.; Wilson, J.; Millman, K. J.; Mayorov, N.; Nelson, A. R. J.; Jones, E.; Kern, R.; Larson, E.; Carey, C. J.; Polat, İ.; Feng, Y.; Moore, E. W.; VanderPlas, J.; Laxalde, D.; Perktold, J.; Cimrman, R.; Henriksen, I.; Quintero, E. A.; Harris, C. R.;

Archibald, A. M.; Ribeiro, A. H.; Pedregosa, F.; van Mulbregt, P. SciPy 1.0: fundamental algorithms for scientific computing in Python. *Nature Methods* **2020**, *17*, 261–272.

(S4) Seabold, S.; Perktold, J. Statsmodels: Econometric and Statistical Modeling with Python. Proceedings of the 9th Python in Science Conference. 2010; pp 92–96.

#### S2. Pattern formation of bovine serum albumin (BSA)

##### Statistical analysis of BSA data

This section reports the full statistical output for the comparison of native and denatured BSA across three lipid chemistries (DOPG, DOPC, and DOTAP). All tests were performed in Python using scipy.stats and statsmodels. Asterisks indicate statistical significance after Holm-Bonferroni correction: *** $p < 0.001$; ** $p < 0.01$; * $p < 0.05$.

**Table ST1.** $I/I_{bulk}$ for native versus denatured BSA at each lipid chemistry: Mann-Whitney $U$ tests with Holm-Bonferroni correction for three pairwise comparisons (one per lipid). An effect size of 1.00 indicates perfect separation between the native and denatured distributions.

| Lipids | Comparison | Median BSA (native) | Median dBSA | Fold difference | *U* | *p*-value |
|---|---|---|---|---|---|---|
| DOPC | BSA vs dBSA | 1.34 | 7.31 | 5.43 | 0 | < 0.001 *** |
| DOPG | BSA vs dBSA | 1.42 | 3.65 | 2.56 | 0 | < 0.001 *** |
| DOTAP | BSA vs dBSA | 1.35 | 9.14 | 6.72 | 0 | < 0.001 *** |

Sample sizes: native BSA $n = 19$ (DOPC), 15 (DOPG), 16 (DOTAP); denatured BSA $n = 21$ (DOPC), 35 (DOPG), 22 (DOTAP). $U = 0$ in every comparison reflects zero overlap between the native and denatured intensity distributions: the largest native value is below the smallest denatured value in each lipid (Reported $p$-values are Holm-corrected).

**Table ST2.** Two-way analysis of variance (ANOVA) on $I/I_{bulk}$ for BSA, with state (native, denatured) and lipid (DOPG, DOPC, DOTAP) as factors. Type II sums of squares; total $n = 128$ droplets; residual degrees of freedom = 122. Partial eta-squared ($\eta^2_p$) reported as effect size.

| Term | Sum of squares | *df* | *F* | *p*-value | $\eta^2_p$ |
|---|---|---|---|---|---|
| State (native, denatured) | 63.78 | 1, 122 | 1533.75 | < 0.001 *** | 0.92 |
| Lipid (DOPG, DOPC, DOTAP) | 6.53 | 2, 122 | 78.50 | < 0.001 *** | 0.56 |
| State × Lipid | 4.49 | 2, 122 | 54.01 | < 0.001 *** | 0.47 |

The significant state × lipid interaction ($\eta^2_p = 0.47$) indicates that the magnitude of the lipid-dependent response differs between native and denatured BSA: native BSA shows only a small, lipid-independent response, whereas denatured BSA shows a strong, lipid-dependent response.

**Table ST3.** Pairwise comparisons of $I/I_{bulk}$ across the three lipid chemistries within denatured BSA: Mann-Whitney *U* tests with Holm-Bonferroni correction for three pairwise comparisons. Effect size is the rank-biserial correlation; sign indicates the direction of the difference. Significance codes: *** $p < 0.001$.

| Protein | Comparison | *U* | *p-value* | effect size |
|---|---|---|---|---|
| dBSA | DOPC vs DOPG | 723 | < 0.001 *** | -0.97 |
| dBSA | DOPC vs DOTAP | 141 | < 0.03 * | 0.39 |
| dBSA | DOPG vs DOTAP | 23 | < 0.001 *** | 0.94 |

Sample sizes: dBSA $n = 21$ (DOPC), 35 (DOPG), 22 (DOTAP). The medians follow the monotonic order DOPG (3.65) < DOPC (7.31) < DOTAP (9.14). All three pairwise comparisons remain significant after the Holm-Bonferroni correction.

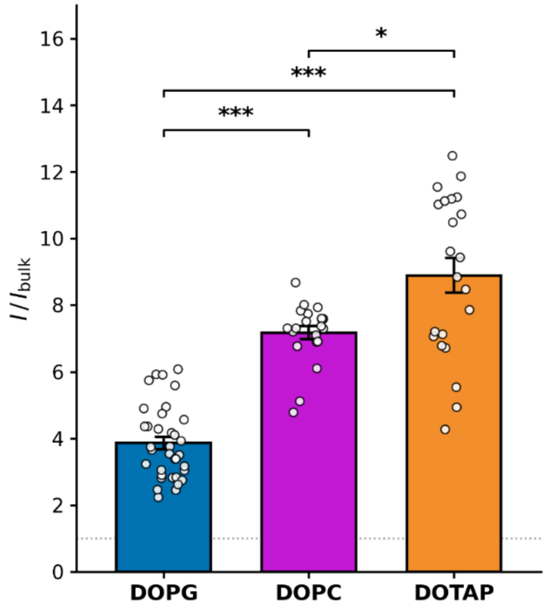


**Figure S1:** Pairwise comparisons of $I/I_{bulk}$ across lipid compositions in denatured BSA, labeled with ThT.

Bar plot showing $I/I_{bulk}$ for dBSA in DOPG (anionic), DOPC (zwitterionic), and DOTAP (cationic) droplets. Bars represent (mean ± SEM); open circles, individual droplets. The dotted reference line at $I/I_{bulk} = 1$ indicates the unconfined baseline. Brackets denote Mann-Whitney U pairwise comparisons with Holm-Bonferroni correction across the three lipid pairs (*** $p < 0.001$; * $p < 0.05$; see Table ST3 for exact values). The medians of $I/I_{bulk}$ fall in the monotonic order DOPG (n = 35, median 3.66) < DOPC (n = 21, median 7.31) < DOTAP (n = 22, median 9.14), consistent with electrostatic favorability for the anionic dBSA.

**Native BSA with different lipid chemistries**

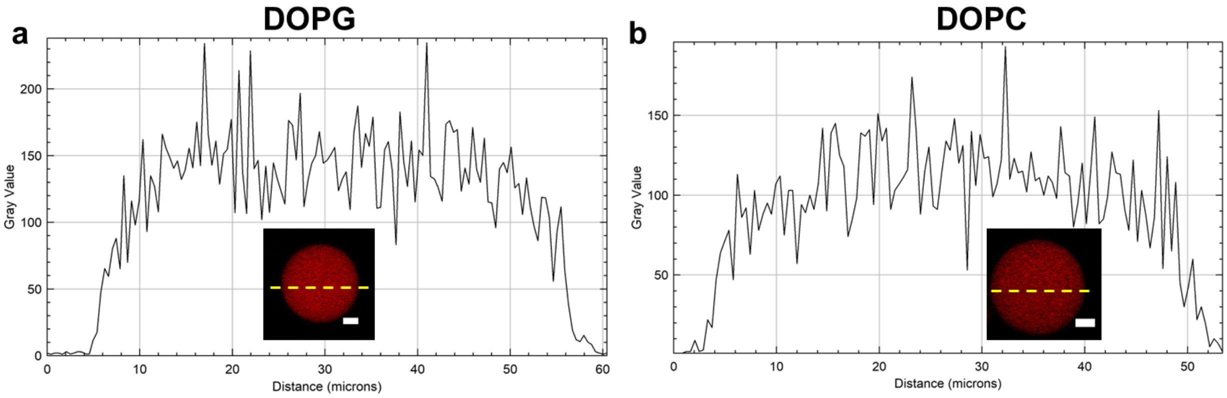


**Figure S2:** Plot profiles of native BSA labeled with Texas Red for lipids (a) DOPG, and (b) DOPC. The scale bar = 10 µm.

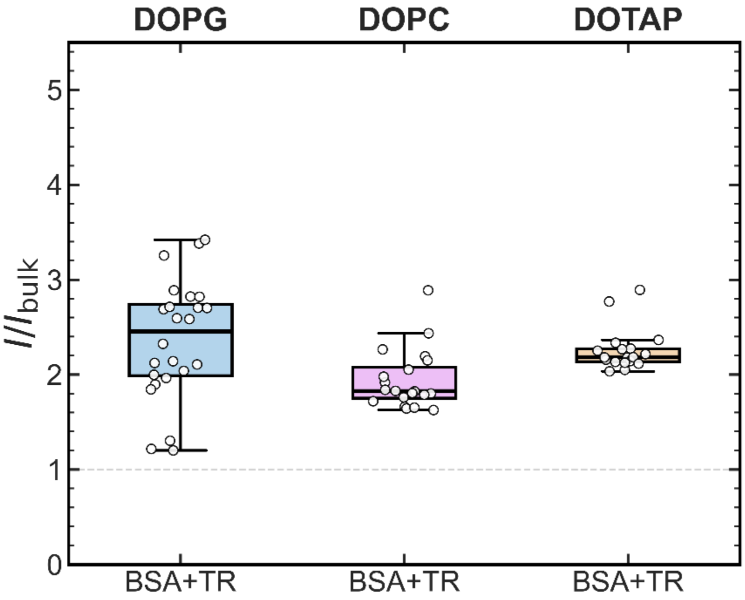


**Figure S3:** Comparisons of $I/I_{bulk}$ across lipid chemistries in native BSA, when labeled with Texas Red.

#### S3. Pattern formation of Lysozyme (Lyz)

##### Statistical analysis of Lyz data

This section reports the full statistical output for the comparison of native and denatured Lyz across three lipid chemistries (DOPG, DOPC, and DOTAP). All tests were performed in Python using scipy.stats and statsmodels. Asterisks indicate statistical significance after Holm-Bonferroni correction: *** $p < 0.001$; ** $p < 0.01$; * $p < 0.05$.

**Table ST4.** Native versus denatured lysozyme at each lipid chemistry: Mann-Whitney *U* tests with Holm-Bonferroni correction for three pairwise comparisons (one per lipid). Effect size is the rank-biserial correlation; positive values indicate higher dLyz medians.

| Lipids | Comparison | Median Lyz (native) | Median dLyz | Fold difference | U | p-value |
|---|---|---|---|---|---|---|
| DOPC | Lyz vs dLyz | 1.16 | 1.47 | 1.26 | 67 | < 0.001 *** |
| DOPG | Lyz vs dLyz | 1.36 | 2.26 | 1.67 | 63 | < 0.001 *** |
| DOTAP | Lyz vs dLyz | 1.45 | 1.63 | 1.12 | 196 | 0.001 ** |

Sample sizes: native Lyz $n$ = 13 (DOPC), 15 (DOPG), 20 (DOTAP); denatured Lyz $n$ = 37 (DOPC), 38 (DOPG), 41 (DOTAP). The largest separation between native and denatured distributions occurs at DOPG, the complementary anionic lipid.

**Table ST5.** Two-way analysis of variance (ANOVA) on $I/I_{\text{bulk}}$ for lysozyme, with state (native, denatured) and lipid (DOPG, DOPC, DOTAP) as factors. Type II sums of squares; total $n$ = 164 droplets; residual degrees of freedom = 158. Partial eta-squared ($\eta^2_p$) reported as effect size.

| **Term** | **Sum of squares** | ***df*** | ***F*** | ***p*-value** | $\eta^2_p$ |
|---|---|---|---|---|---|
| State (native, denatured) | 3.31 | 1, 158 | 60.17 | < 0.001 *** | 0.28 |
| Lipid (DOPG, DOPC, DOTAP) | 4.65 | 2, 158 | 42.26 | < 0.001 *** | 0.35 |
| State × Lipid | 0.71 | 2, 158 | 6.49 | 0.002 ** | 0.08 |

The state × lipid interaction is significant ($\eta^2_p$ = 0.08) but of substantially smaller magnitude than for BSA ($\eta^2_p$ = 0.46; Table ST2), indicating that the change in the lipid response between native and denatured states is smaller for lysozyme than for BSA.

**Table ST6.** Pairwise comparisons of $I/I_{\text{bulk}}$ across the three lipid chemistries within denatured lysozyme: Mann-Whitney *U* tests with Holm-Bonferroni correction for three pairwise comparisons. Effect size is the rank-biserial correlation; the sign indicates the direction of the difference. Significance codes: *** $p < 0.001$; ** $p < 0.01$.

| protein | comparison | *U* | *p-value* | effect size |
|---|---|---|---|---|
| dLyz | DOPC vs DOPG | 168 | < 0.001 *** | 0.76 |
| dLyz | DOPC vs DOTAP | 460 | 0.002 * | 0.39 |
| dLyz | DOPG vs DOTAP | 1300 | < 0.001 *** | -0.67 |

Sample sizes: dLyz $n = 37$ (DOPC), 38 (DOPG), 41 (DOTAP). The $I/I_{\text{bulk}}$ of DOPG-containing comparisons are separated strongly from the other lipid (DOPC vs DOPG and DOPG vs DOTAP, both $p < 0.001$), while DOPC and DOTAP differ at a smaller effect size ($p = 0.003$).

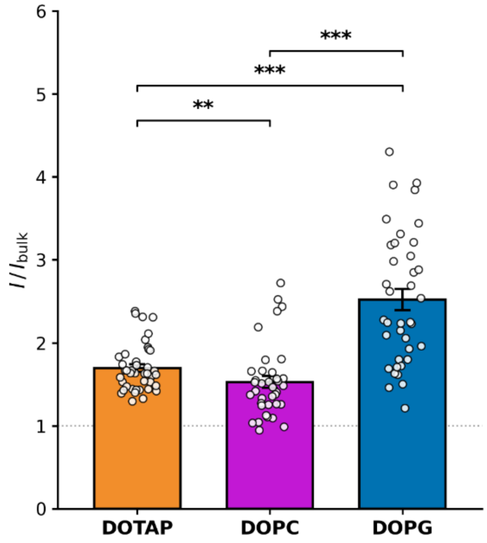


**Figure S4:** Pairwise comparisons of $I/I_{bulk}$ across lipid chemistries in denatured Lyz, labeled with ThT.

Bar plot showing $I/I_{\text{bulk}}$ for dLyz in DOPG (anionic), DOPC (zwitterionic), and DOTAP (cationic) droplets. Bars represent (mean ± SEM); open circles, individual droplets. The dotted reference line at $I/I_{\text{bulk}} = 1$ indicates the unconfined baseline. Brackets denote Mann-Whitney U pairwise comparisons with Holm-Bonferroni correction across the three lipid pairs (*** $p < 0.001$; * $p < 0.05$; see Table ST6 for exact values). The medians of $I/I_{\text{bulk}}$ follow the order DOPC (1.47) < DOTAP (1.63) < DOPG (2.26), with DOPG separating from the two non-complementary lipids.

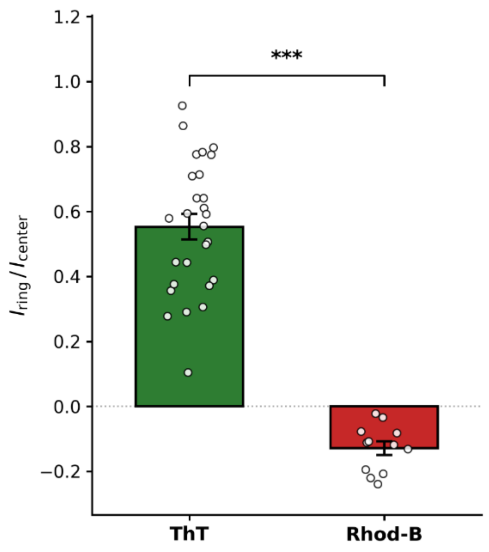


**Figure S5:** Total protein remains uniformly distributed in dLyz/DOPG droplets (visualized using Rhod-B), while β-sheet structure localizes at the lipid interface (visualized using ThT).

Bar plot of the normalized rim-enhancement ratio $I_{\text{ring}}$ / $I_{\text{center}}$, measured in dLyz/DOPG droplets. The ThT (green), reporting β-sheet structure; and Rhod-B (red), reporting total protein density. ThT fluorescence shows strong positive rim enhancement (median +0.58, IQR 0.38–0.71, $n$ = 27 droplets), while protein dye fluorescence (Rhod-B) shows a slightly negative rim ratio (median −0.11, IQR −0.20 to −0.08, $n$ = 12 droplets). The two distributions did not overlap (Mann-Whitney $U$ test, $p < 0.001$ ***, with effective size = −1.00). Bars represent mean ± SEM; open circles, individual droplets; dotted reference line at $I_{\text{ring}}$ / $I_{\text{center}}$, = 0 indicates the no-rim baseline.

**Native Lyz with different lipid chemistries**

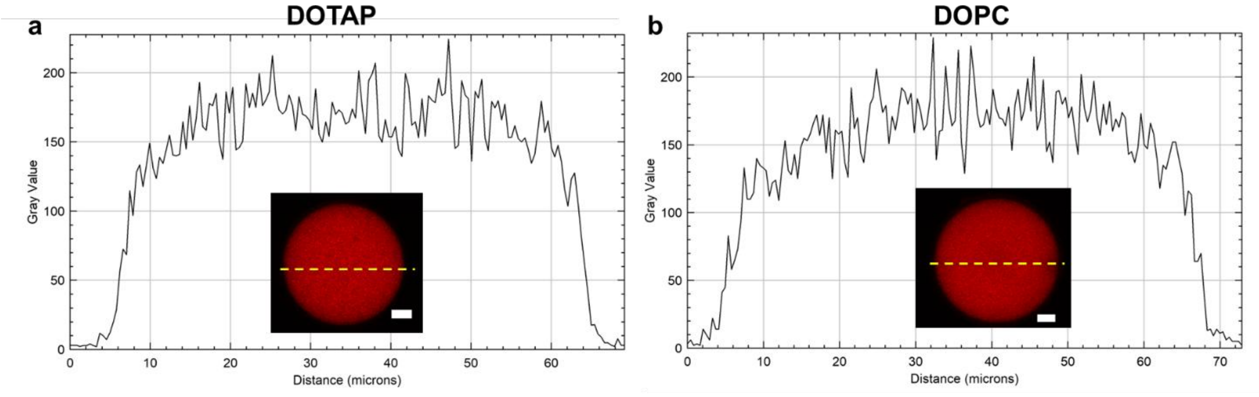


**Figure S6:** Fluorescence intensity profiles of dye-labeled native lysozyme along the droplet center for lipids (a) DOPG, and (b) DOPC. An example of the droplet is illustrated for each case. The scale bar = 10 μm

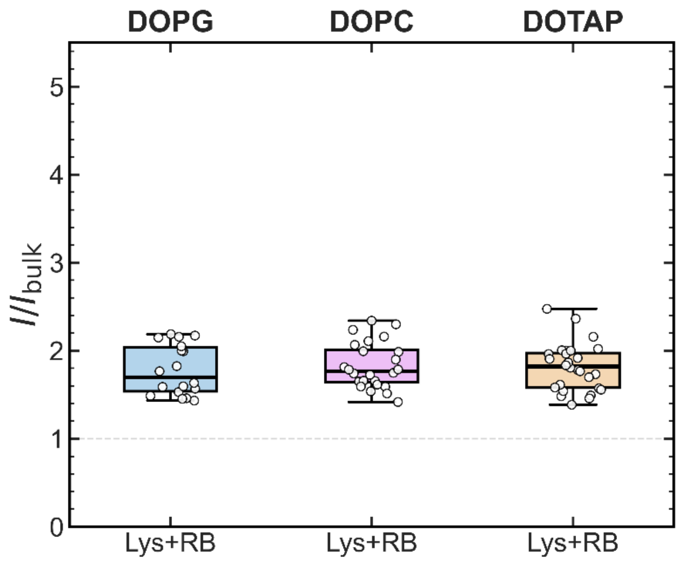


**Figure S7:** Comparisons of $I/I_{bulk}$ across lipid chemistries for RB-labeled native lysozyme (Lyz) in droplets. From left to right: DOPG, DOPC, and DOTAP.

**Comparison of dLyz and dBSA with cross lipid chemistry**

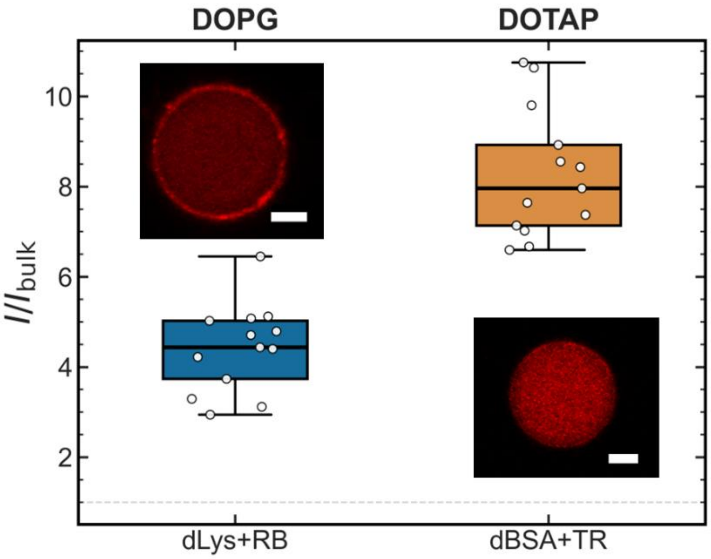


**Figure S8:** Comparisons of $I/I_{bulk}$ across lipid chemistries, (left) dye-labelled dLyz inside DOPG droplets (P(+)L(-)) and dye-labelled dBSA insie DOTAP droplets (P(-)L(+)). An example of the droplet is shown for each case. The scale bar = 10 μm.

**Homologs of Lyz and BSA with cross lipid chemistry**

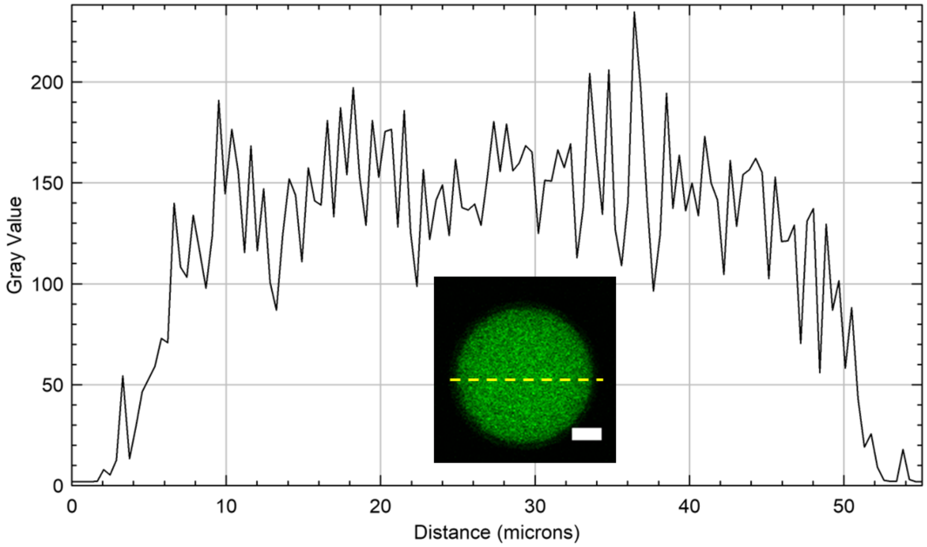


**Figure S9:** ThT fluorescence intensity profiles of denatured HSA in DOTAP droplets along the droplet center. The scale bar in the confocal image is 10 µm.

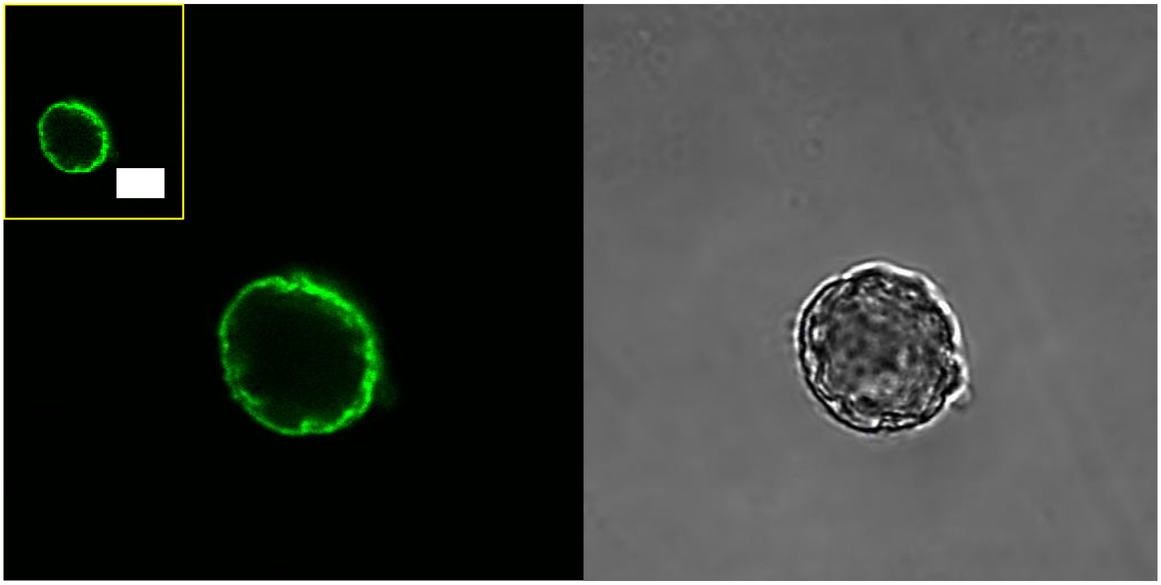

**Figure S10:** Denatured human lysozyme labeled with ThT in DOPG-coated droplets, showing buckling in a droplet with a radius of approximately 7 µm (inset). Enlarged views of the confocal and bright-field images are shown. Scale bars: 10 µm.
